# Kinetic hierarchy of Kai protein complex formation governs the cyanobacterial circadian oscillator

**DOI:** 10.64898/2026.03.06.710049

**Authors:** Ken Morishima, Yasuhiro Yunoki, Tatsuro Oda, Koichi Mayumi, Rintaro Inoue, Masaaki Sugiyama

**Affiliations:** Institute for Integrated Radiation and Nuclear Science, Kyoto University, 2-1010 Asashironishi, Kumatori, Sennan-gun, Osaka, 590-0494, Japan; The Institute for Solid State Physics, The University of Tokyo, 5-1-5 Kashiwanoha, Kashiwa, Chiba, 277-8581, Japan

## Abstract

Cyanobacterial circadian oscillations arise from a phosphorylation cycle of KaiC that is coupled to reversible complex formation with KaiA and KaiB. Although representative assemblies such as A_2_C_6_, B_6_C_6_, and A_12_B_6_C_6_ have been structurally characterized, quantitative understanding of their stoichiometry, affinity, and formation kinetics during the oscillation cycle remains limited. Here, we systematically quantified AC, BC and ABC complex formation using phosphorylation-mimetic KaiC and controlled protein mixing ratios by integrating analytical ultracentrifugation with small-angle X-ray/neutron scattering.

A_2_C_6_ formation exhibits graded dependence on the phosphorylation state of KaiC and occurs rapidly in solution. In contrast, B_6_C_6_ formation behaves in a switch-like manner, showing strong selectivity for the hyperphosphorylation mimic and proceeding on a slow timescale (∼ 6 h). Upon B_6_C_6_ formation, KaiA rapidly associates with the complex to generate A_*n*_B_6_C_6_, with KaiA occupancy *n* determined by the mixing ratio through fast redistribution among coexisting complexes. This behavior enables dynamic allocation of KaiA among clock complexes.

Together, these quantitative insights delineate a hierarchy of assembly dynamics—fast, graded AC exchange; slow, state-selective BC formation; and rapid KaiA redistribution—revealing a mechanistic basis for the dynamic regulation of the Kai oscillator.

## INTRODUCTION

Circadian clocks generate ∼24 h rhythms that coordinate physiology across diverse organisms. Cyanobacteria offer a uniquely minimal and tractable system: robust circadian phosphorylation–dephosphorylation oscillations of KaiC can be reconstituted in vitro using only KaiA, KaiB and KaiC in the presence of ATP^1-5^.

In the canonical view, KaiC cycles through ordered phosphorylation states at S431 and T432^6,7^, and this cycle is coupled to reversible complex formation. KaiA–KaiC (AC) complexes promote phosphorylation, whereas KaiB–KaiC (BC) and KaiA–KaiB–KaiC (ABC) complexes are associated with dephosphorylation^8-11^. Structural analyses have established representative assemblies such as A_2_C_6_, B_6_C_6_, and A_12_B_6_C_6_, defining the architectures through which Kai proteins interact^8,10,12-31^.

However, the oscillator operates far from equilibrium, in which key state parameters—including the phosphorylation state of KaiC and the effective concentrations of Kai proteins—vary continuously during the oscillation cycle. Consequently, complex formation occurs under continuously evolving conditions rather than fixed biochemical states. Understanding the system therefore requires determining which complexes form under these evolving conditions, how rapidly they assemble, and which association reactions dominate the overall dynamics of the oscillator.

Here we address this problem by analyzing phosphorylation-mimetic KaiC under controlled mixing ratios and directly quantifying complex formation with a unified, label-free combination of analytical ultracentrifugation (AUC) and small-angle X-ray/neutron scattering (SAXS/SANS). This framework reveals distinct formation behaviors of AC, BC and ABC complexes, uncovering a hierarchy of complex formation that regulates the dynamics of the Kai oscillator.

## RESULTS

### Formation behavior of AC complex

To reveal the affinity, stoichiometry, and kinetics of AC complex formation, we first analyzed binary mixtures of KaiA and KaiC phosphorylation-mimic mutants (AA, AE, DE, and DA, mimicking S/T, S/*p*T, *p*S/*p*T, and *p*S/T, respectively; see METHODS) by combining AUC and time-resolved SAXS. Figure 1**a** shows AUC-derived weight concentration distributions from titration of KaiA against KaiC_AA_ (hypophosphorylation mimic) with fixed concentration (2.0 mg/mL) as a representative example (see Supplementary Figure 1 for other mutants). The molar mixing ratio per a protomer was defined as [KaiA]: [KaiC] = *x*: 6, and measurements were performed one hour after mixing. With increasing *x*, the peak corresponding to the C_6_ (*s*_20,w_ = 11.2 S) shifted continuously toward higher *s*_20,w_-values. At sufficiently high KaiA mixing ratios (*x* > 6), the peak converged at *s*_20,w_ = 12.3 S (Fig. 1**b**), and an additional peak corresponding to free A_2_ appeared at *s*_20,w_ = 3.6 S. The molecular mass calculated at *s*_20,w_ = 12.3 S was 417 kDa, which agreed with that expected for A_2_C_6_ (Supplementary Note 1 and Supplementary Table 1). The same result was obtained for the other KaiC mutants, therefore, the stoichiometry of A_2_ binding to C_6_ does not exceed 1:1 irrespective of KaiC phosphorylation state or KaiA mixing ratio.

**Figure 1.**
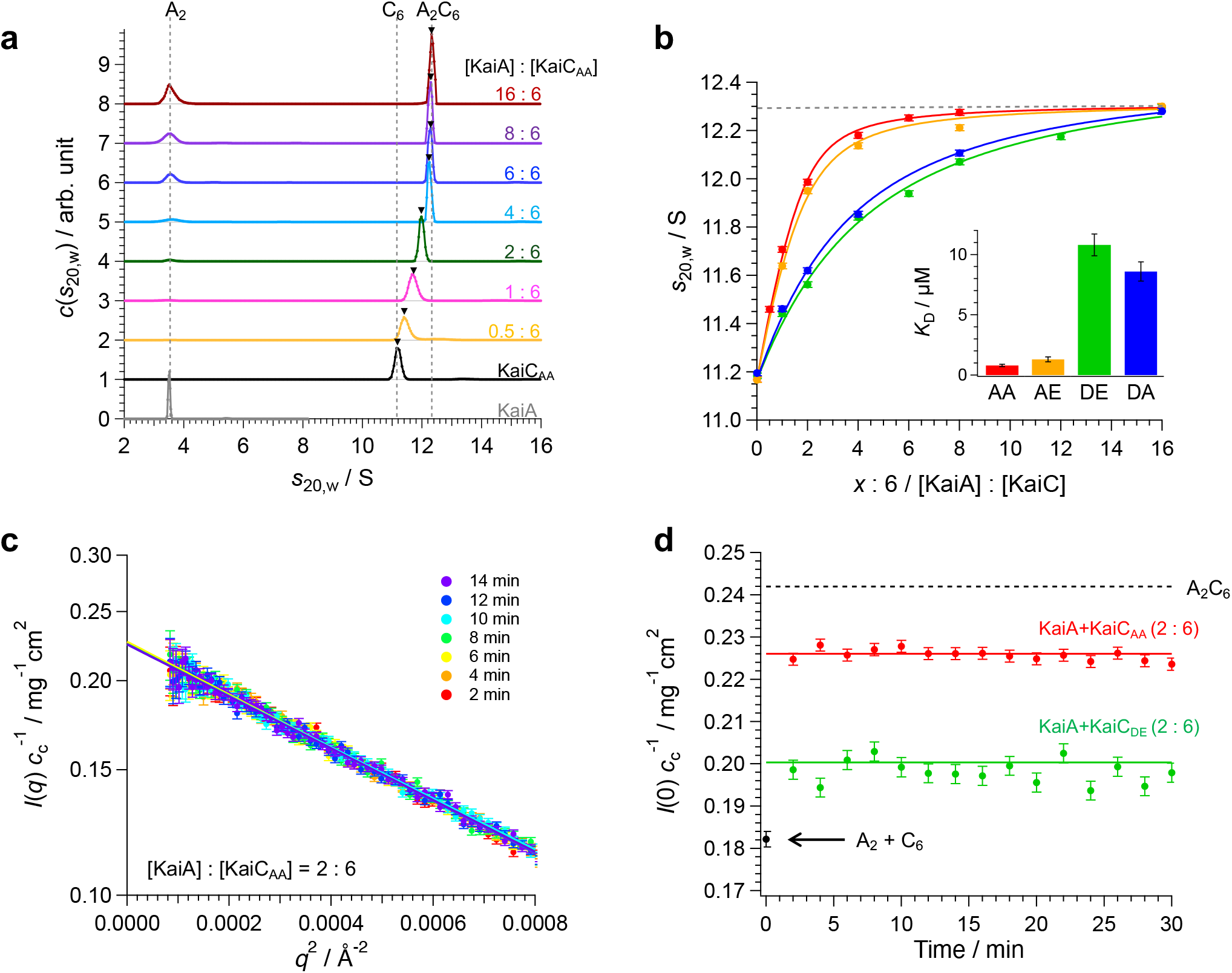
Association-dissociation behavior of AC complex. **a**. Weight-concentration distributions *c*(*s*_20,w_) offered by AUC for KaiA, KaiC_AA_, and mix solutions of KaiA+KaiC_AA_ with various mixing ratios *x* of KaiA to KaiC; [KaiA]: [KaiC] = *x*: 6 (*x* = 0 – 16). The concentration of KaiC was fixed at 2.0 mg/mL. Black triangles mean the peak position of C_6_/A_2_C_6_ component. **b**. Mixing ratio-dependence of the *s*_20,w_-value of the peak corresponding to C_6_/A_2_C_6_ component. Red, yellow, green, and blue circles represent the result for KaiC_AA_, KaiC_AE_, KaiC_DE_, and KaiC_DA_, respectively. Solid lines are the fitting curve which gives the average sedimentation coefficient *s*_av_ for C_6_/A_2_C_6_ component (Supplementary Note 2). Broken line corresponds to the *s*_20,w_-value of A_2_C_6_ (*s*_20,w,AC_). Inset represents *K*_D_-value depending on the KaiC-mutant. **c**. Guinier plot of the time resolved-SAXS profile for the KaiA+KaiC_AA_ solution at *x* = 2. Solid lines mean square fitting lines with Guinier formula. **d**. Red and green circles represent the time dependences of forward scattering intensity normalized by KaiC-concentration (*I*(0)*c*_c_^-1^) for KaiA+KaiC_AA_ and KaiA+KaiC_DE_ solutions at *x* = 2, respectively. Solid red and green lines mean the calculated value based on *K*_D_ for each mutant (Table 1 and Supplementary Note 3). Black circle is the values for the dissociated (A_2_ and C_6_) state which calculated by the summation of the experimental value for solo A_2_ and C_6_ (Supplementary Figure 2 and Supplementary Table 1). Broken black line represents the calculated values for fully associated (A_2_C_6_) states.

At low KaiA mixing ratios (*x* ≤ 6), a single peak was observed between 11.2 S (= C_6_) and 12.3 S (= A_2_C_6_). Considering that A_2_ and C_6_ are the fundamental unit in solution, this peak is attributed to contributions from both C_6_ and A_2_C_6_, appearing as a single merged peak whose sedimentation coefficient reflects their population ratio. Such merged peaks are typically observed when sedimentation coefficients of components are inseparably close or when the association-dissociation between components is fast relative to the experimental timescale^32^. Based on the result of kinetics described below, this case is attributed to the latter scenario.

Assuming an equilibrium of A_2_ + C_6_ ⇄A_2_C_6_, the weight-averaged sedimentation coefficient *s*_av_ was expressed as the following function with a dissociation constant *K*_D_ as the sole adjustable parameter^33^ (see detail in Supplementary Note 2).

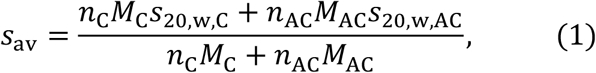

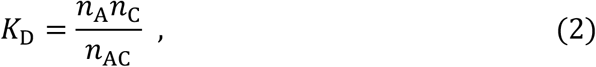

where *n*_*i*_, *M*_*i*_, *s*_20,w,*i*_ (*i* = C or AC) are number concentration, molecular mass, and sedimentation coefficient, respectively, for C_6_ and A_2_C_6_. Using this expression for *s*_av_ with the optimal *K*_D_-value (Inset of Fig. 1**b** and Supplementary Table 2), the mixing-ratio dependence of the experimental *s*_20,w_-values was reproduced (solid line in Fig. 1**b**). This good fitting strongly supports the interpretation that the association-dissociation behavior is reasonably described by the simple equilibrium A_2_ + C_6_ ⇄A_2_C_6_, and further demonstrates that, complexes such as A_4_C_6_ and A_6_C_6_ are not present in solution.

The *K*_D_-values offered by the fitting analysis followed the order AA < AE << DA < DE (Figure 1**b** and Supplementary Table 2), indicating stronger affinity between A_2_ and C_6_ at lower phosphorylation states. Notably, the phosphorylation state of S431 strongly influences AC complex formation and the difference in *K*_D_ between AA and DE mutants was approximately 14-fold. This value is substantially smaller than the 100–1000-fold difference assumed in the previous mathematical model^34^, indicating that modulation of A_2_ and C_6_ affinity across the phosphorylation cycle is less drastic than previously predicted. This result suggests that the effect of the phosphorylation-state dependence of the KaiA–KaiC affinity to the mechanism of the circadian oscillation may be smaller than previously assumed. We next examined the kinetics of AC complex formation using time-resolved SAXS. To measure the AC complex formation speed, SAXS profiles were collected every 2 min immediately after mixing KaiA and KaiC at a mixing ratio of *x* (= 6[KaiA]/[KaiC]) = 2 (Fig. 1**c**). The scattering profiles *I*(*q*) at each time point (*q*: magnitude of the scattering vector) were fitted with Guinier approximation^35^ (Supplementary Note 3), yielding the forward scattering intensity *I*(0) and gyration radius *R*_g_. Figure 1**d** shows the time courses of *I*(0) obtained by time-resolved SAXS at two-minute intervals after mixing KaiA and KaiC (AA and DE as red and green circles, respectively). The *I*(0) values corresponding to the fully dissociated state (A_2_ + C_6_) and the fully associated state (A_2_C_6_) were calculated (refer to Supplementary Note 3, Supplementary Figure 2, and Supplementary Table 3), and are represented in Fig. 1**d** by a black circle and dashed line, respectively. For both phosphorylation-state mutants, *I*(0) already exceeded that of the fully dissociated state at two minutes after mixing and subsequently reached a constant value. In addition, the *R*_g_ also remained constant, indicating no time-dependent structural change (Supplementary Figure 3). Using the *K*_D_ values obtained from AUC (Supplementary Table 2), the number concentrations of A_2_, C_6_, and A_2_C_6_ were calculated with Eqs. S8–S10 in Supplementary Note 2 (*n*_A_: *n*_C_: *n*_AC_ = 0.35: 0.35: 0.65 for AA and 0.61: 0.61: 0.39 for DE under the mixing ratio of *x* = 2), and the corresponding *I*(0) values were estimated using Eq. S18 in Supplementary Note 3. The calculated values (solid red and green lines for AA and DE in Fig. 1**d**) agreed well with the experimental data. These results indicate a rapid association-dissociation equilibrium between A_2_ and C_6_ that is established within at most two minutes. This behavior is consistent with previous observations by high-speed AFM^13^ and further demonstrates that it can be captured in a label-free and non-perturbative manner in solution by SAXS.

### Formation behavior of BC complex

To serve as a basis for subsequent analysis of the ABC complex, we next examined BC complex formation using binary mixtures of KaiB and KaiC mutants by combining AUC, SANS, and time-resolved SAXS. Figure 2**a** exhibits AUC-derived weight concentration distributions from titration of KaiB against KaiC_DE_ (hyperphosphorylation mimic) with fixed concentration (1.0 mg/mL) as a representative example (see Supplementary Figure 4 for other mutants). The molar mixing ratio per a protomer was defined as [KaiB]: [KaiC] = *y*: 6, and measurements were conducted following 24 h incubation at 30 °C after mixing. With increasing *y*, the peak corresponding to C_6_ (*s*_20,w_ = 11.2 S) shifted continuously toward higher *s*_20,w_-values and converged at *s*_20,w_ = 12.3 S in *y* ≥ 6 (Fig. 2**b**). The molecular mass calculated at *s*_20,w_ = 12.3 S was 415 kDa, which agreed with that expected for B_6_C_6_ (Supplementary Note 1 and Supplementary Table 1).

**Figure 2.**
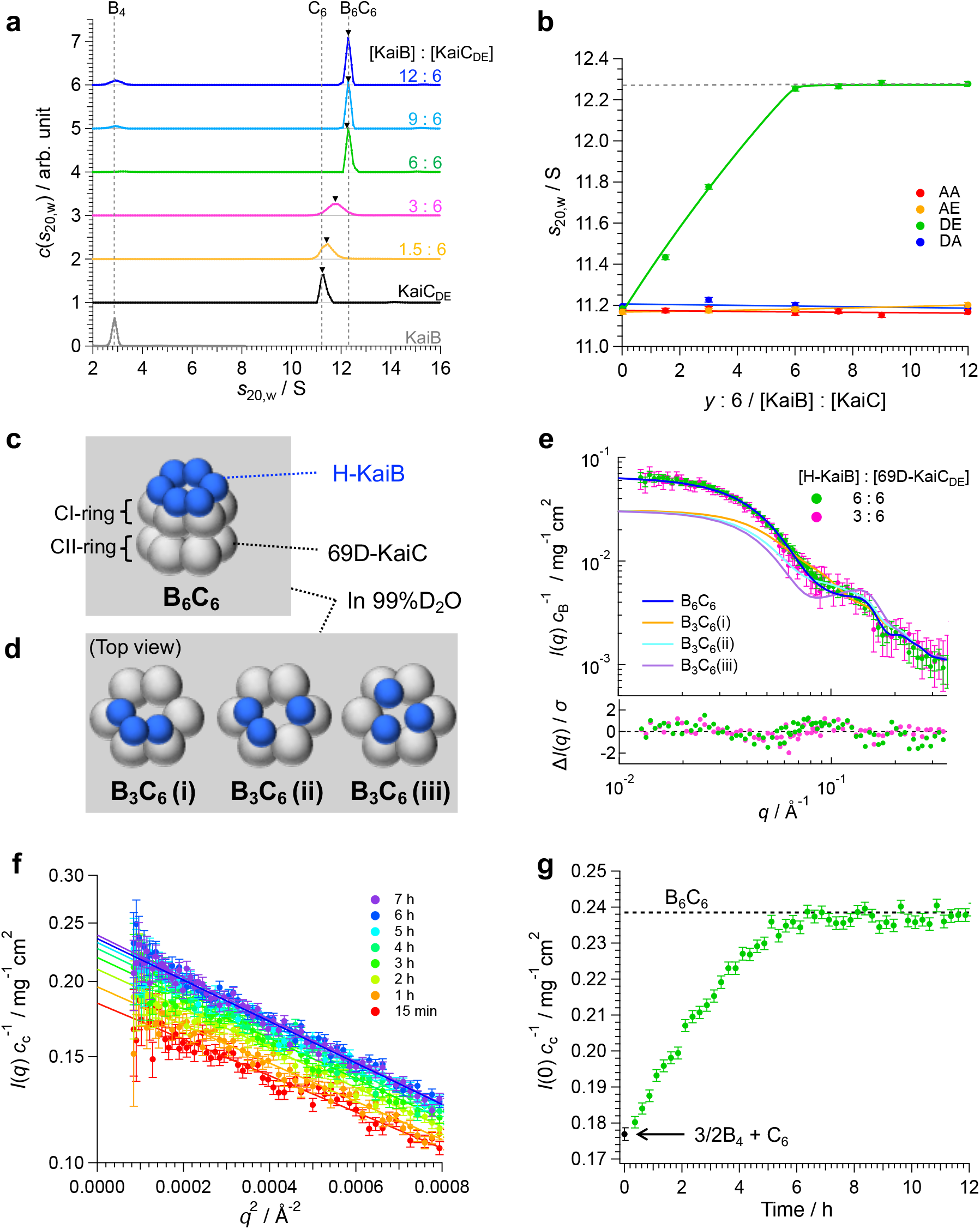
Association-dissociation behavior of BC complex. **a**. Weight-concentration distributions *c*(*s*_20,w_) offered by AUC for KaiB solution, KaiC_DE_ solution and mix solution of KaiB+KaiC_DE_ with various mixing ratios of KaiB to KaiC_DE_, [KaiB]: [KaiC_DE_] = *y*: 6 (*y* = 0 – 12). The concentration of KaiC was fixed at 1.0 mg/mL. Black triangles mean the peak position of C_6_/B_6_C_6_ component. **b**. The mixing ratio-dependence of the *s*_20,w_-value of the peak corresponding to C_6_/B_6_C_6_ component. Red, yellow, green, and blue circles represent the result for KaiC_AA_, KaiC_AE_, KaiC_DE_, and KaiC_DA_, respectively. Solid lines are the fitting curve which gives the average sedimentation coefficient *s*_av_ for C_6_ and B_6_C_6_. (Supplementary Note 2). Broken line corresponds to the *s*_20,w_-value of B_6_C_6_. **c**. Schematic illustration of B_6_C_6_ complex composed of H-KaiB and 69D-KaiC in 99%D_2_O. **d**. Schematic illustration of B_3_C_6_ complexes with three different KaiB configurations from top view (CI-ring side). **e**. The iCM-SANS profiles for the mix solutions of H-KaiB and 69D-KaiC_DE_ in 99%D_2_O buffer at *y* = 3 (magenta circles) and *y* = 6 (green circles) of mixing ratios. Scattering intensities were normalized by the weight concentration of KaiB, *c*_B_. Solid blue, yellow, cyan, and purple lines represent the calculated scattering intensity for B_6_C_6_, B_3_C_6_(i), B_3_C_6_(ii), and B_3_C_6_(iii), respectively, composed of H-KaiB and 69D-KaiC_DE_ in 99%D_2_O based on the reference structure (refer to METHOD). Lower panel show the residuals between the experimental scattering profiles (magenta: *y* = 3 and green: *y* = 6) and calculated scattering profile for B_6_C_6_. Here, *σ* means the standard error of experimental scattering intensities. **f**. Guinier plot for time resolved-SAXS profile for the KaiB+KaiC_DE_ solution at *y* = 6. Solid lines mean square fitting lines with Guinier formula. **g**. Green circles represent the time dependence of forward scattering intensity normalized by *c*_c_ for the KaiB+KaiC_DE_ solutions. Black circle is the values for the dissociated (B_4_ and C_6_) state which calculated by the summation of the experimental value for solo B_6_ and C_6_ (Supplementary Figure 2 and Supplementary Table 1). Broken black line represents the calculated values for fully associated (B_6_C_6_) states.

At low mixing ratios (*y* < 6), only a single peak between C_6_ and B_6_C_6_ was observed. The origin of this single peak can be attributed either to insufficient peak-separation of the C_6_ and B_6_C_6_ components, leading to a merged peak, or to the formation of intermediate stoichiometry complexes B_*m*_C_6_. To resolve this ambiguity, we conducted inverse contrast-matching SANS (iCM-SANS)^21,36,37^ for mix solution of hydrogenated KaiB (H-KaiB) and 69% deuterated KaiC_DE_ (69D-KaiC_DE_) in 99% D_2_O. Under this condition, C_6_ is effectively contrast-matched and therefore scattering-invisible (Supplementary Figure 5), allowing selective observation of B_*m*_ within B_*m*_C_6_ (Fig. 2**c** and **d**). The normalized iCM-SANS profile *I*(*q*)*c*_B_^-1^, where *c*_B_ is the weight concentration of H-KaiB, for mixtures of H-KaiB and 69D-KaiC_DE_ at *y* = 3 (magenta circles) and 6 (green circles) overlapped well with each other (Fig.2**e**). These scattering profiles, as well as the corresponding forward scattering intensities *I*(0)*c*_B_^-1^ and gyration radii *R*_g_, agreed well with the calculated values for the B_6_-ring within B_6_C_6_ (blue solid line in Fig. 2**e** and Supplementary Table 4). If KaiB protomer were evenly distributed among all C_6_ hexamers at *y* = 3, B_3_C_6_ would be expected as the dominant binding ratio. However, none of the possible configurations of B_3_C_6_ as displayed in Fig. 2**d** reproduced the experimental scattering profile at *y* =3 (yellow, cyan, and purple solid lines in Fig. 2**e** and Supplementary Table 4). Therefore, KaiB predominantly exists as B_6_C_6_ even at *y* = 3 as observed at *y* = 6, indicating highly cooperative binding of KaiB to C_6_, which results in a uniquely stable stoichiometric assembly B_6_C_6_.

Based on these observations, only C_6_ and B_6_C_6_ coexist in solution even at non-saturating ratios (*y* < 6), giving rise to a single AUC peak whose sedimentation coefficient reflects their relative populations (Fig. 2**b**). The observed *s*_20,w_-values were well reproduced by the population-weighted average sedimentation coefficient *s*_av_ of C_6_ and B_6_C_6_ calculated from 3/2B_4_+C_6_ ⇄B_6_C_6_ association-dissociation equilibrium (Fig. 2**b** and Supplementary Note 2). Here, the green solid line in Fig. 2**b** shows the titration curve calculated with *K*_D_ = 10^-3^ μM. Although smaller *K*_D_-values cannot be reliably determined within the concentration range accessible to AUC, the KaiB–KaiC_DE_ interaction was estimated to have *K*_D_ < 10^-3^ μM (Supplementary Figure 6), demonstrating that it is substantially stronger than the KaiA–KaiC interaction.

Notably, no BC complex formation was detected for phosphorylation-mimicking mutants other than KaiC_DE_, indicating that the ability to form the BC complex differs sharply between the hyperphosphorylated state (DE) and all other states.

We next investigated the formation kinetics of the BC complex using time-resolved SAXS. Figure 2**f** shows the time course of *I*(*q*)*c*_c_^-1^, where *c*_c_ is the weight concentration of KaiC, obtained by time-resolved SAXS measurements immediately after mixing KaiB and KaiC_DE_ at a mixing ratio of *y* = 6. The *I*(0)*c*_c_^-1^-value initially corresponded to that of the dissociated state (B_4_ and C_6_) and gradually increased with time, reaching the value corresponding to that of B_6_C_6_ after approximately six hours (Fig. 2**g**). This slow formation is consistent with previous observations of delayed KaiB–KaiC association reported using EPR spectroscopy and other spectroscopic approaches^18,19^. Importantly, we quantitatively resolve multiple defining features of BC complex formation within a single experimental framework, using non-perturbative solution measurements, simultaneously demonstrating that B_6_C_6_ assembly is highly cooperative, strongly selective for the hyperphosphorylated state, and proceeds on a timescale of hours under defined biochemical conditions.

### Formation behavior of ABC complex

We next investigated how KaiA associates with the preformed B_6_C_6_ complex to generate the ABC complex regarding stoichiometry and kinetics. AUC measurements were performed on samples titrating KaiA into preformed B_6_C_6_ (prepared using KaiC_DE_; see METHODS), with B_6_C_6_ kept at a constant concentration (1.0 mg/mL) (Fig. 3**a**). The molar mixing ratio was defined as [KaiA]: [B_6_C_6_] = *z*: 1, and measurements were conducted one hour after mixing. With increasing *z*, the main peak shifted continuously from *s*_20,w_ = 12.3 S, corresponding to B_6_C_6_, toward higher *s*_20,w_-values, and reached *s*_20,w_ = 17.4 S at *z* = 24 (triangles in Fig. 3**a** and Supplementary Figure 7). The molecular mass calculated at *s*_20,w_ = 17.4 S was 811 kDa, which corresponds to that of A_12_B_6_C_6_ (Supplementary Note 1 and Supplementary Table 1).

**Figure 3.**
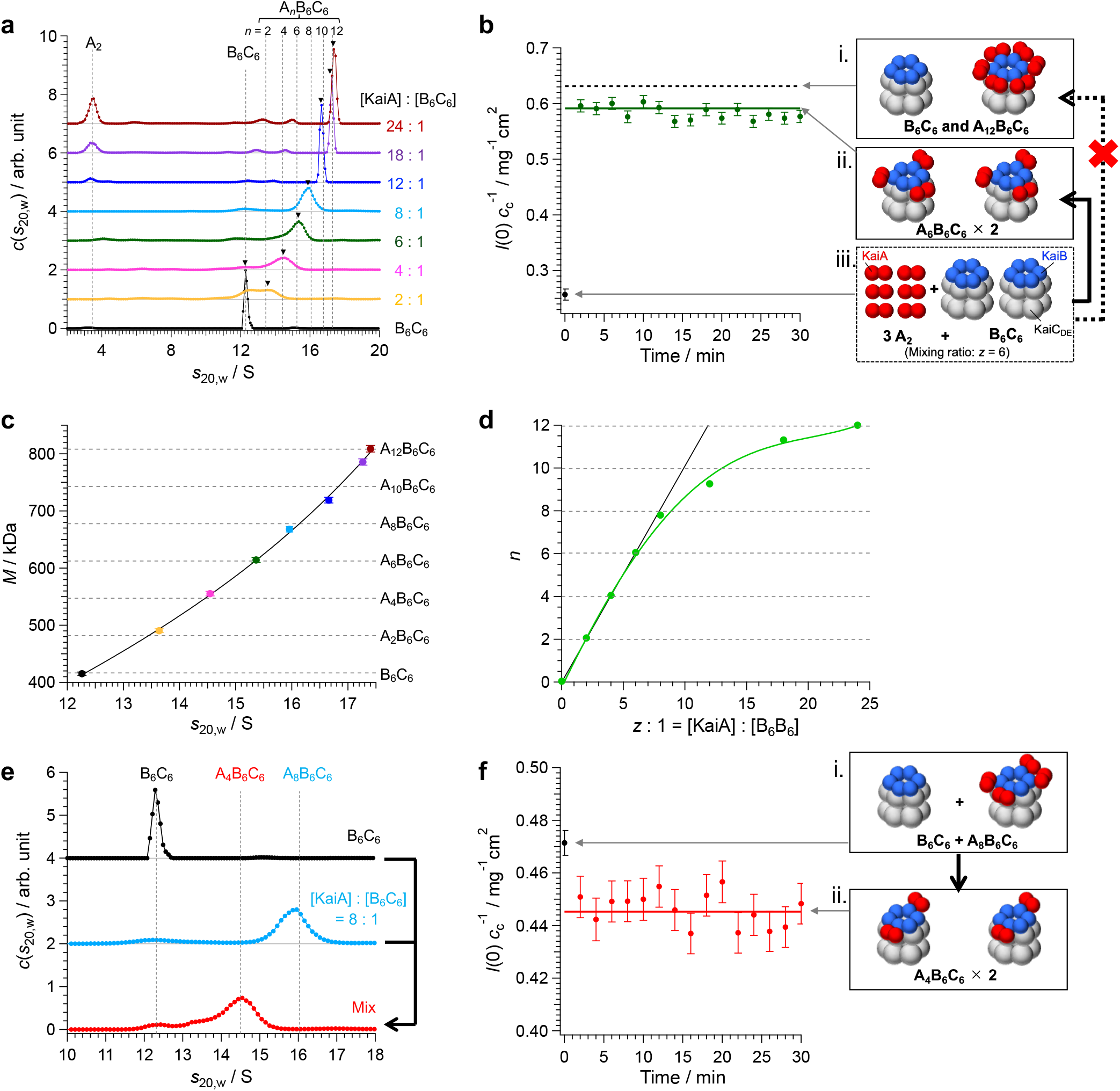
Association-dissociation behavior of ABC complex. **a**. Weight-concentration distributions *c*(*s*_20,w_) offered by AUC for mix solution of KaiA+B_6_C_6_ with various mixing ratios of KaiA to B_6_C_6_, [KaiA]: [B_6_C_6_] = *z*: 1 (*z* = 0 – 24). The concentration of B_6_C_6_ was fixed at 1.0 mg/mL. Black triangles mean the peak position of A_*n*_B_6_C_6_ component. **b**. Green circles represent the forward scattering intensity normalized by KaiC-concentration (*I*(0)*c*_c_^-1^) provided by the time-resolved SAXS after mixing KaiA and B_6_C_6_ (Supplementary Figure 8**a**). Solid green line corresponds to the calculated *I*(0)*c*_c_^-1^-value for A_6_B_6_C_6_. Broken black line corresponds to that for the summation of B_6_C_6_ and A_12_B_6_C_6_, respectively. **c**. Relationship between *s*_20,w_ and molecular mass *M* of A_*n*_B_6_C_6_. The molecular mass was derived from peak *s*_20,w_-value and *f*/*f*_0_ (Supplementary Figure 7**b** and Supplementary Note 1). The color of circles corresponds to the mixing ratio shown in panel **a**. Solid line shows the fitting curve with Eq.S19 in Supplementary Note 4. Broken lines represent the molecular masses of A_*n*_B_6_C_6_ (*n* = 0, 2, 4, 6, 8, 10, and 12) calculated from amino acid sequences. **d**. Relationship between mixing ratio *z* and association number *n* of KaiA in A_*n*_B_6_C_6_. Solid green line is the fitting line with polynomial function for eye-guide. Solid black line represents *n* = *z*. **e**. Black, cyan, and red circles show the *c*(*s*_20,w_) for B_6_C_6_, A_8_B_6_C_6_, and their mixture, respectively. Broken lines represent *s*_20,w_-values for B_6_C_6_, A_4_B_6_C_6_, and A_8_B_6_C_6_. **f**. Red circles are forward scattering intensity normalized by KaiC-concentration (*I*(0)*c*_c_^-1^) provided by the time-resolved SAXS after mixing B_6_C_6_ and A_8_B_6_C_6_ (Supplementary Figure 8**a**). Solid red line represents *I*(0)*c*_c_^-1^ calculated for A_4_B_6_C_6_. Black circle represents the summation of the experimental value for B_6_C_6_ and A_8_B_6_C_6_.

The peaks observed between 12.3 S (B_6_C_6_) and 17.4 S (A_12_B_6_C_6_) at intermediate mixing ratios (0 < *z* < 24) could arise either from population-dependent merging of B_6_C_6_ and A_12_B_6_C_6_ or from the formation of intermediate A_*n*_B_6_C_6_ (0 < *n* < 12). To distinguish between these possibilities, we performed time-resolved SAXS measurements after mixing KaiA and B_6_C_6_ at *z* = 6 as a representative condition (Supplementary Figure 8**a**). At this mixing ratio of *z* = 6, if A_12_B_6_C_6_ were the only stable ABC complex, coexistence of B_6_C_6_ and A_12_B_6_C_6_ would be expected at a 1:1 molar ratio (Fig. 3**b**-i). In contrast, uniform distribution of KaiA among all B_6_C_6_ would yield A_6_B_6_C_6_ as the dominant complex In contrast, uniform distribution of KaiA among all B_6_C_6_ would yield A_6_B_6_C_6_ as the dominant complex (Fig. 3**b**-ii). The forward scattering intensity obtained by SAXS reached, within two minutes after mixing, a value that agreed with the calculated value for A_6_B_6_C_6_ (dark green line in Fig. 3**b**). This result indicates that A_12_B_6_C_6_ is not uniquely stable in solution; rather, distinct A_*n*_B_6_C_6_ are stabilized according to the mixing ratio.

The relationship between the *s*_20,w_-values (Fig. 3**a**, triangles) and the corresponding molecular mass of A_*n*_B_6_C_6_ was derived as Fig. 3**c** (refer to Supplementary Note 1 and Supplementary Figure 7). Figure 3**d** represents the relationship between the mixing ratio *z* (= [KaiA]/[B_6_C_6_]) and the association number *n* of KaiA within A_*n*_B_6_C_6_. At *z* ≤ 8, *n* = *z* holds (solid black line in Fig. 3d), indicating that all added KaiA binds B_6_C_6_ to form ABC complex. In contrast, at *z* > 8, A_*n*_B_6_C_6_ with *n* < *z* form, and free A_2_ appears in the AUC profiles (Fig. 3**a**). These results indicate that KaiA binding to B_6_C_6_ becomes progressively weaker as *n* increases, such that higher-order KaiA association occurs with lower affinity. Taken together with the SAXS results (Fig. 3**b**), these findings suggest that, when KaiA and B_6_C_6_ coexist at any mixing ratio, KaiA is evenly distributed among B_6_C_6_ such that A_*n*_B_6_C_6_ with the smallest possible *n* consistent with the mixing ratio is rapidly formed.

When the mixing ratio *z* is changed, KaiA is expected to undergo dissociation and reassociation from preformed A_*n*_B_6_C_6_, such that the even-distribution rule of KaiA is re-established and A_*n*_’ B_6_C_6_ with a new association number *n*’ is formed according to the new mixing ratio. To access this scenario, we mixed A_8_B_6_C_6_ (light blue circles; a KaiA+B_6_C_6_ mix solution at *z* = 8) with an equal amount of B_6_C_6_ (black circles) and analyzed the distribution of A_*n*_B_6_C_6_ by AUC (Fig. 3**e**). As predicted from the KaiA mixing ratio (*z* = 4) after combining the two solutions, A_4_B_6_C_6_ became the dominant complex in the mixture (red circles in Fig. 3**e**). Time-resolved SAXS measurements for the same sample immediately after mixing (Supplementary Figure 8**b**) revealed that the forward scattering intensity rapidly (within at most two minutes) changed from the value corresponding to the sum of A_8_B_6_C_6_ and B_6_C_6_ (black circles in Fig. 3**f**) to that corresponding to A_4_B_6_C_6_ (red line in Fig. 3**f**), reaching equilibrium. Collectively, these results demonstrate that KaiA within the ABC complex obeys an even-distribution rule and that A_*n*_B_6_C_6_ with an association number *n* of KaiA appropriate for the mixing ratio is formed rapidly.

## DISCUSSION

This study provides a quantitative, solution-based comparison of AC, BC and ABC complex formation across defined phosphorylation-mimetic states and controlled mixing ratios. The use of phosphorylation-mimetic KaiC mutants provides an experimental framework for systematic analysis of complex formation across defined conditions by eliminating phosphorylation heterogeneity and time-dependent state changes during the oscillation cycle. This design allowed stoichiometry, apparent affinities, and formation kinetics to be quantitatively assigned to well-defined biochemical states.

A central finding is the qualitative contrast between AC and BC complexes in their dependence on the phosphorylation state of KaiC. AC complex formation showed a relatively graded behavior: A_2_C_6_ forms across all phosphorylation-mimetic states with continuously varying binding affinity. In contrast, BC complex formation exhibited a much sharper dependence, with stable B_6_C_6_ detected only for the hyperphosphorylation mimic (DE) and absent for the other states. These observations indicate that BC complex formation behaves in a more switch-like manner than AC formation with respect to KaiC phosphorylation state.

The phosphorylation-state dependence of A_2_C_6_ formation quantified here was substantially smaller than the 100–1000-fold affinity difference assumed in the allosteric model proposed by van Zon et al^34^. Instead, we observed an approximately 14-fold difference between hypophosphorylated and hyperphosphorylated mimics. Although this modulation of KaiA–KaiC affinity may still contribute to oscillator behavior, the magnitude observed here suggests that additional mechanisms may participate in generating robust circadian oscillations. Our results point to BC complex formation as a potential contributor in this regard. In particular, B_6_C_6_ formation shows both a much stronger phosphorylation-state dependence and a markedly slower formation timescale than A_2_C_6_. These features raise the possibility that, during the oscillatory cycle, the conversion of hyperphosphorylated KaiC to B_6_C_6_ could represent a rate-limiting step that contributes to synchronization mechanism^34,38^ of phosphorylation phases across the KaiC population.

We also found that once B_6_C_6_ forms, KaiA associates rapidly to form A_*n*_B_6_C_6_. Importantly, KaiA binding affinity decreases as the association number *n* increases, leading to rapid redistribution of KaiA among coexisting B_6_C_6_ and A_*n*_B_6_C_6_ complexes. Consequently, A_*n*_B_6_C_6_ with the smallest *n* compatible with the mixing ratio are preferentially formed. This redistribution behavior indicates that KaiA occupancy within A_*n*_B_6_C_6_ is dynamically determined by the balance between affinity and protein stoichiometry.

From a kinetic perspective, KaiA association and dissociation within both A_2_C_6_ and A_*n*_B_6_C_6_ occur much faster than the formation of B_6_C_6_. Therefore, when these complexes coexist during the oscillatory process^39^, KaiA is expected to exchange between them on a substantially shorter timescale. Such rapid exchange would enable dynamic allocation of KaiA according to relative affinities and component concentrations, allowing the system to redistribute KaiA efficiently as the phosphorylation state of KaiC evolves.

Overall, our results suggest that Kai-system complex formation is governed by an interplay between (i) graded modulation of AC affinity, (ii) phosphorylation-state-selective and slow BC formation, and (iii) rapid KaiA redistribution within A_2_C_6_ and A_*n*_B_6_C_6_. This integrated picture provides experimentally based constraints for refining mechanistic models of circadian oscillation in the Kai-system. More broadly, these findings illustrate how dynamic protein complex formation with distinct kinetic and stoichiometric properties can generate robust biochemical oscillations.

## ACKNOWLEDGEMENTS

We are grateful to Prof. Koichi Kato, Dr. Hirokazu Yagi (ExCELLS/IMS/Nagoya City University), and Prof. Kazuki Terauchi (Ritsumeikan University) for providing samples and useful advice. We are also grateful to Dr. Masahiro Shimizu, Mr. Ritsuki Sakamoto, Mr. Yuki Katsuno (Kyoto University) for their assistance of experiments and discussion. SANS measurements were conducted with the approval of the Institute for Solid State Physics (ISSP), The University of Tokyo and the Japan Atomic Energy Agency (JAEA) under proposal Nos. 22914, 23556, 24404, 24536, and 24540. AUC and SAXS measurements were conducted at KURNS under proposal Nos. R3069, R4010, R5016, R6076, and R7089.

## DATA AVAILABILITY

All data in this article are available from the authors upon reasonable request.

## METHODS

### Expression and purification of proteins

KaiA, KaiB and KaiC from *Synechococcus elongatus* PCC 7942 were expressed in *Escherichia coli* BL21DE3 codon plus (KaiA and KaiB) and DH5α (KaiC). To prepare hydrogenated KaiA, KaiB and KaiC, the bacterial cells were grown in terrific broth medium. To prepare partially deuterated KaiC for iCM-SANS, the bacterial cells were grown in M9 minimal H_2_O/D_2_O mixture media containing hydrogenated and deuterated glucose (1,2,3,4,5,6,6-*d*7, and 98%; Cambridge Isotope Laboratories, Inc., Tewksbury, MA, USA) as previously described^40^. KaiA was generated as a hexahistidine (His)-tagged recombinant protein and purified after cleavage of the His-tag as previously described^26,41^. KaiB was generated as a glutathione S-transferase (GST)-tagged recombinant protein and purified after the cleavage of the GST-tag as previously described^24^. KaiC was generated as a Strep-tagged recombinant protein and was purified as previously described^42^. In this study, the following phosphorylation mimic KaiC mutants were utilized; S/T–mimicking S431A/T432A mutant (KaiC_AA_), S/*p*T–mimicking S431A/T432E mutant (KaiC_AE_), *p*S/*p*T–mimicking S431D/T432E mutant (KaiC_DE_), and *p*S/T–mimicking S431D/T432A mutant (KaiC_DA_).

All proteins were finally purified by size exclusion chromatography (SEC) with an eluent buffer containing 50 mM sodium phosphate (pH 7.8), 150 mM sodium chloride, 5 mM magnesium chloride, 0.5 mM EDTA, 1 mM dithiothreitol, 1 mM adenosine triphosphate (ATP), 50 mM glutamic acid, and 50 mM arginine. The eluent buffer was prepared in D_2_O for iCM-SANS measurement and in H_2_O for other measurements.

The deuteration degree of KaiC_DE_ and D_2_O ratio in solvent of iCM-SANS sample, determined using mass spectrometry and infrared spectroscopy according to previously reported methods^40^, were 69% and 99%, respectively.

### Sample preparation

KaiA+KaiC titration sample series was prepared by mixing KaiA and KaiC solutions with the composition of [KaiA]: [KaiC] = *x*: 6 (*x* = 0 – 16), where [KaiA] and [KaiC] mean the molar concentration for their monomer. The partial concentration of KaiC was fixed at 2.0 mg/mL. The solution was subjected to AUC measurements after incubation at 30 °C for one hour.

KaiB+KaiC titration sample series was prepared by mixing KaiB and KaiC solutions with the composition of [KaiB]: [KaiC] = *y*: 6 (*y* = 0 – 12), where [KaiB] and [KaiC] mean the molar concentration for their monomer. The partial concentration of KaiC was fixed at 1.0 mg/mL. The solution was subjected to AUC measurements after incubation at 30 °C for 24 hours.

KaiA + B_6_C_6_ titration sample series was prepared by mixing KaiA and B_6_C_6_ solutions with the composition of [KaiA]: [B_6_C_6_] = *z*: 1 (*z* = 0 – 24). Here, B_6_C_6_ was prepared by the size exclusion chromatography (SEC) of KaiB + KaiC_DE_ mix solution ([KaiB]: [KaiC_DE_] = 6: 6) after incubation at 30 °C for 24 hours. The partial concentration of B_6_C_6_ was fixed at 1.0 mg/mL. The KaiA+B_6_C_6_ solution was subjected to AUC measurements after incubation at 30 °C for one hour.

### Analytical ultracentrifugation (AUC)

AUC measurements were performed using ProteomeLab XL-I (Beckman Coulter Inc., Brea, CA, USA). The samples were loaded into cells equipped with 12 mm optical path aluminums center piece. Sedimentation velocity analysis was performed using Rayleigh interference optics at 30 °C. The rotor speed was set at 60,000 rpm and the measurement was carried out for one hour for all measurements. The weight-concentration distributions as a function of sedimentation coefficient *c*(*s*_20,w_) and average frictional ratio *f/f*_0_ were obtained through computational fitting to the time evolution of sedimentation data with the multi-component Lamm formula using SEDFIT software version 15.01c^43^. The sedimentation coefficient was normalized to be the value at 20 °C in pure water, *s*_20,w_^44^.

### Small-angle X-ray scattering (SAXS)

SAXS measurements were performed using a laboratory-based instrument NANOPIX (Rigaku, Japan) equipped with a high-brilliance point-focused generator of a Cu-Kα source MicroMAX-007 HFMR (Rigaku, Japan) (wavelength = 1.54 Å). Scattered X-rays were measured using a HyPix-6000 Hybrid Photon Counting detector (Rigaku, Japan) composed of 765 × 813 pixels with the spatial resolution of 100 μm. For all samples, the sample-to-detector distance was set to be 1330 mm, with which the covered *q*-range was 0.01 Å^-1^ ≤ *q* ≤ 0.20 Å^-1^ (*q*: magnitude of scattering vector). Two-dimensional scattering patterns were converted to one-dimensional scattering profiles using SAngler software^45^. After correction by the transmittance and subtraction of buffer scattering, the absolute scattering intensity was obtained using the standard scattering intensity of water (1.632 × 10^-2^ cm^-1^)^46^. All measurements were performed at 30 °C.

### Small-angle neutron scattering (SANS)

SANS measurements were performed with SANS-U^47,48^ at JRR-3 (Japan Atomic Energy Agency, JAEA, Ibaraki, Japan). A neutron beam with a wavelength (λ) of 6.0Å and its distribution (Δλ/λ) of 10% was irradiated to the samples. The diameter of the beam was set to be 15 mm. The source-to-sample distance (boron-coated collimator length) was set to 4,000 mm. The sample-to-detector distances were set to 4,000 and 1,030 mm, covering the magnitude of scattering vector *q* range of 0.012 – 0.3 Å^−1^. Scattered neutrons were recorded using a two-dimensional detector (ORDELA, Oak Ridge, TN, USA). Two-dimensional scattering patterns were converted to one-dimensional scattering profiles using Red2D software (https://github.com/hurxl/Red2D). After correction by the transmittance and subtraction of buffer scattering, the scattering intensity was converted to the absolute one using the standard scattering intensity of H_2_O (0.89 cm^−1^)^49^. All measurements were performed at 30 °C.

AUC-SAS treatment was applied to eliminate the influence of aggregated molecules from the experimental SANS profile. Details of the AUC-SAS treatment have been described elsewhere^50,51^.

Scattering curve of BC complex were calculated based on the reference structure (PDB: code 5n8y^27^) using CRYSON^52^.

