## supplementally Information for "Kinetic hierarchy of Kai protein complex formation governs the cyanobacterial circadian oscillator"

**The file includes:**

- Supplementary Note 1-4 p. 2-6
- Supplementary Figures 1-8 p. 7-14
- Supplementary Tables 1-4 p. 15
- References p. 16

**Supplementary Note:**

**Supplementary Note 1. Relationship of sedimentation coefficient and molecular mass:**

From the balance of the centrifugal force, buoyant force, and viscous drag acting on a particle with molecular mass $M$ in a centrifugal field, the following equation is obtained.

$$s = \frac{M(1-\rho\bar{v})}{N_A f}, \qquad \text{(S1)}$$

where $s$, $\rho$, $\bar{v}$, $N_A$, and $f$ are sedimentation coefficient, density of solvent, partial specific volume of the particle, Avogadro's number, and frictional coefficient, respectively. We introduce the friction coefficient $f_0$ of a hypothetical spherical particle with the same molecular mass and partial specific volume as the sedimenting particle. Based on Stokes' law, the friction coefficients $f$ and $f_0$ are expressed using the solvent viscosity $\eta$.

$$f = 6\pi\eta R_H,\ \ f_0 = 6\pi\eta R_{H0} \qquad \text{(S2)}$$

Here, $R_H$ and $R_{H0}$ represent the hydrodynamic radii of the sedimenting and hypothetical spherical particles, respectively, and $R_{H0}$ is given by

$$R_{H0} = \left(\frac{3M\bar{v}}{4\pi N_A}\right)^{\frac{1}{3}}. \qquad \text{(S3)}$$

From Eqs.S1-S3, we obtained the relationship of the molecular mass and sedimentation coefficient with frictional ratio $f/f_0$ as follows:

$$M = \left[\frac{6\pi\eta N_A}{(1-\rho\bar{v})}\left(\frac{3\bar{v}}{4\pi N_A}\right)^{\frac{1}{3}}\left(\frac{f}{f_0}\right)\right]^{\frac{3}{2}} s^{\frac{3}{2}}\ . \qquad \text{(S4)}$$

**Supplementary Note 2. Sedimentation coefficient of merged peak from $C_6/A_2C_6$ and $C_6/B_6C_6$ components:**

**(i) Weight-averaged sedimentation coefficient $s_{av}$ for $C_6/A_2C_6$ component**

Assuming an equilibrium of $A_2+C_6 \rightleftarrows A_2C_6$, the weight-averaged sedimentation coefficient $s_{av}$ was expressed as follows[1].

$$s_{\mathrm{av}} = \frac{n_{\mathrm{C}} M_{\mathrm{C}} s_{20,\mathrm{w,C}} + n_{\mathrm{AC}} M_{\mathrm{AC}} s_{20,\mathrm{w,AC}}}{n_{\mathrm{C}} M_{\mathrm{C}} + n_{\mathrm{AC}} M_{\mathrm{AC}}}, \qquad \text{(S5)}$$

where $n_i$, $M_i$, $s_{20,w,i}$ ($i$ = C or AC) are number concentration, molecular mass, and sedimentation coefficient, respectively, for $C_6$ and $A_2C_6$. According to the mass action law (Eq.S6), the number concentration of $A_2$, $C_6$, and $A_2C_6$ under the equilibrium are expressed as Eqs.S7-S10.

$$K_{\mathrm{D}} = \frac{n_{\mathrm{A}} n_{\mathrm{C}}}{n_{\mathrm{AC}}}\ . \qquad \text{(S6)}$$

$$n_{\mathrm{AC}} = n_{\mathrm{C,0}} - n_{\mathrm{C}}\ . \qquad \text{(S7)}$$

$$n_{\mathrm{A}} = (x/2) n_{\mathrm{C,0}} - n_{\mathrm{AC}}\ . \qquad \text{(S8)}$$

$$n_{\mathrm{C}} = \frac{1}{2}\left[-A \pm \left(A^2 + 4K_{\mathrm{D}} n_{\mathrm{C,0}}\right)^{0.5}\right]. \qquad \text{(S9)}$$

$$A = K_{\mathrm{D}} + (x/2) n_{\mathrm{C,0}} - n_{\mathrm{C,0}}\ . \qquad \text{(S10)}$$

Here, $K_D$, $n_{\mathrm{A}}$, $n_{\mathrm{C,0}}$ are dissociation constant, number concentration of $A_2$, and preparation number concentration of $C_6$, respectively. Substituting Eqs. S7–S10 into Eq.S5, $s_{av}$ is expressed as a function of $x$ with $K_D$ as the sole adjustable parameter.

**(ii) Weight-averaged sedimentation coefficient $s_{av}$ for $C_6/B_6C_6$ component**

Assuming an equilibrium of $3/2B_4+C_6 \rightleftarrows B_6C_6$, the $s_{\mathrm{av}}$ for $C_6$ and $B_6C_6$ is expressed as follows.

$$s_{\mathrm{av}} = \frac{n_{\mathrm{C}} M_{\mathrm{C}} s_{20,\mathrm{w,C}} + n_{\mathrm{BC}} M_{\mathrm{BC}} s_{20,\mathrm{w,BC}}}{n_{\mathrm{C}} M_{\mathrm{C}} + n_{\mathrm{BC}} M_{\mathrm{BC}}}, \qquad \text{(S11)}$$

Here, $n_i$, $M_i$, $s_{20,w,i}$ ($i$ = C or BC) are number concentration, molecular mass, and sedimentation coefficient, respectively, for $C_6$ and $B_6C_6$. According to the mass action law (Eq. S12), the number concentrations of $B_4$, $C_6$ and $B_6C_6$ are related as Eqs.S13-S16.

$$K_{\mathrm{D}} = \frac{(2/3) n_{\mathrm{B}} n_{\mathrm{C}}}{n_{\mathrm{BC}}}\ . \qquad \text{(S12)}$$

$$n_{\mathrm{BC}} = n_{\mathrm{C,0}} - n_{\mathrm{C}}\ . \qquad \text{(S13)}$$

$$n_{\mathrm{B}} = (3/2)\left[(y/6) n_{\mathrm{C,0}} - n_{\mathrm{BC}}\right]. \qquad \text{(S14)}$$

$$n_{\mathrm{C}} = \frac{1}{2}\left[-B \pm \left(B^2 + 4K_{\mathrm{D}} n_{\mathrm{C,0}}\right)^{0.5}\right]. \qquad \text{(S15)}$$

$$B = K_{\mathrm{D}} + (y/6) n_{\mathrm{C,0}} - n_{\mathrm{C,0}}\ . \qquad \text{(S16)}$$

Here, $n_{\mathrm{B}}$, $n_{\mathrm{C,0}}$ are dissociation constant, number concentration of $B_4$, and preparation number concentration of $C_6$, respectively. Substituting Eqs. S13–S16 into Eq.S11, $s_{\mathrm{av}}$ is expressed as a function of $y$ with $K_{\mathrm{D}}$ as the sole adjustable parameter.

### Supplementary Note 3. Guinier approximation and forward scattering intensity:

The scattering profiles $I(q)$ at each time point ($q$: magnitude of the scattering vector) were fitted with Guinier formula[2] (Eq. S17) in the $q$-range of $q \le 1.3/R_g$.

$$I(q) = I(0)\exp\left(-\frac{R_g^2}{3}q^2\right). \tag{S17}$$

Here, $I(0)$ and $R_g$ are forward scattering intensity and gyration radius, respectively. For a multi-component solution, $I(0)$ is proportional to the second moment of the molecular mass distribution of all components in solution as follows.

$$I(0) = A\sum_i n_i M_i^2 , \tag{S18}$$

where $n_i$ and $M_i$ denote the number concentration and molecular mass of component $i$, respectively. Here, $A \equiv \Delta\rho^2\bar{v}^2/N_A$ ($N_A$: Avogadro's number), assuming that the scattering contrast $\Delta\rho$ and specific volume $\bar{v}$ are identical for all components.

**Supplementary Note 4. Relationship between the molecular mass, sedimentation coefficient, and frictional ratio of $A_nB_6C_6$.**

Since the frictional ratio $f/f_0$ obtained from AUC varies with mixing ratio, reflecting shape changes of $A_nB_6C_6$ as a function of $n$. We accordingly rewrote Eq.1 as follows using $f(s_{20,\mathrm{w}})$ which is the $s_{20,\mathrm{w}}$-dependence of $f/f_0$ (Supplementary Fig. 4**b**).

$$M = a\left[f(s_{20,\mathrm{w}}) \cdot s_{20,\mathrm{w}}\right]^{1.5}, \qquad (S19)$$

where $a$ is a constant. Solid curve in Fig.3**c** represents the fitting curve by Eq.S19, deriving the relationship between $M$ and $s_{20,\mathrm{w}}$ for $A_nB_6C_6$.

### Supplementary Figures:

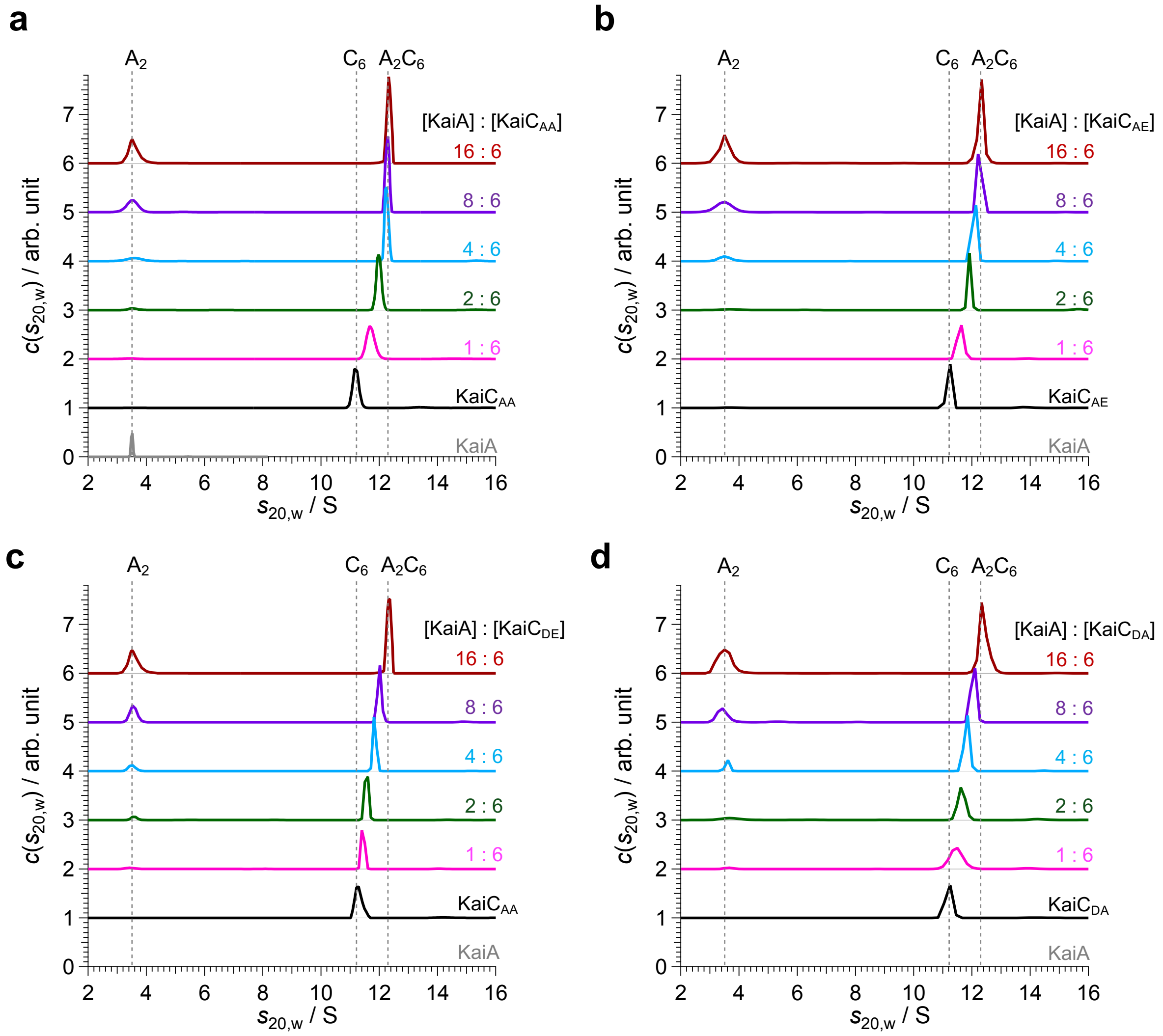


**Supplementary Figure 1. AUC-derived weight concentration distributions from titration of KaiA against KaiC.** Weight-concentration distributions $c(s_{20,\mathrm{w}})$ obtained by AUC for KaiA, KaiC, and mix solutions of KaiA+KaiC with various molar mixing ratios. Concentration of KaiC was fixed at 2.0 mg/mL. Panels **a**, **b**, **c**, and **d** show the results for KaiC$_{AA}$, KaiC$_{AE}$, KaiC$_{DE}$, and KaiC$_{DE}$, respectively.

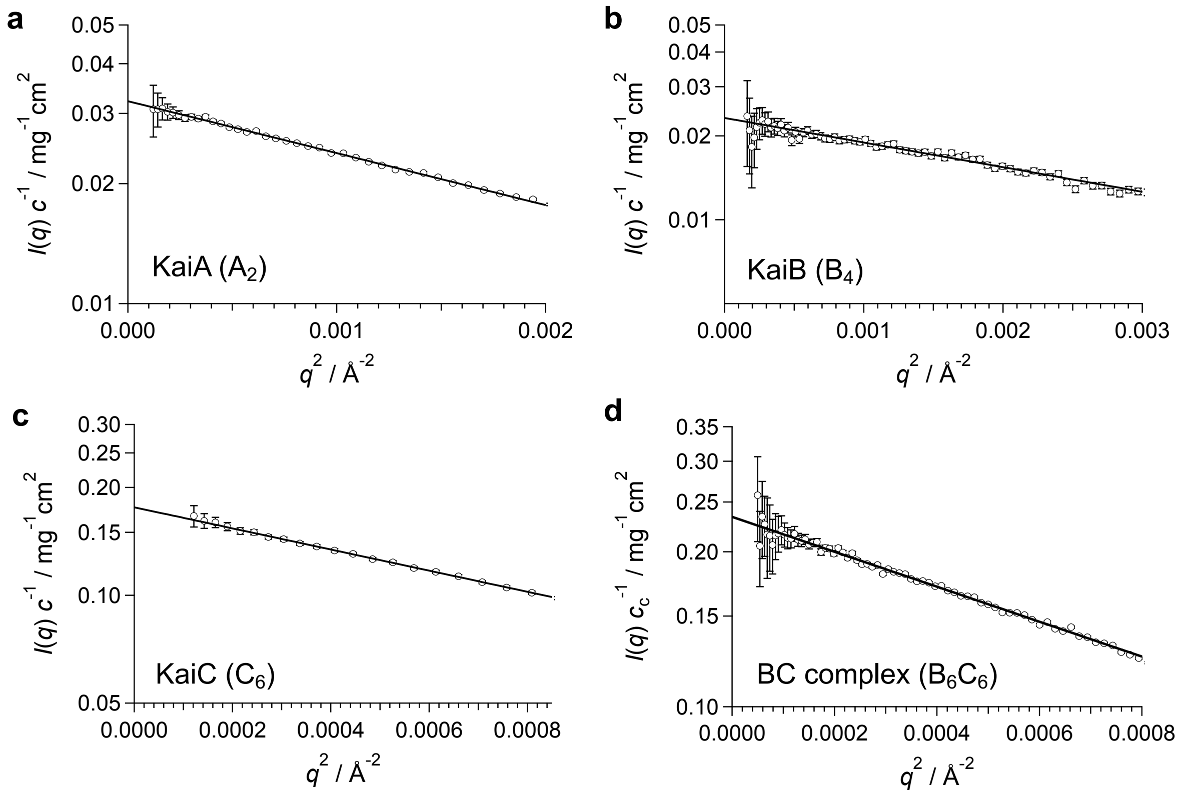


**Supplementary Figure 2. Static SAXS measurement for KaiA, KaiB, KaiC, and BC complex solutions.** Guinier plots of SAXS profiles for **a.** KaiA ($A_2$), **b.** KaiB ($B_4$), **c.** KaiC ($C_6$) and **d.** BC complex ($B_6C_6$). Scattering intensities for KaiA, KaiB, and KaiC are normalized by their weight concentration $c$. Scattering intensities for BC complex are normalized by their weight concentration of KaiC $c_c$. Solid lines represent the least square fitting with Guinier formula. Forward scattering intensities and gyration radii obtained by Guinier analysis were summarized in Supplementary Table 1.

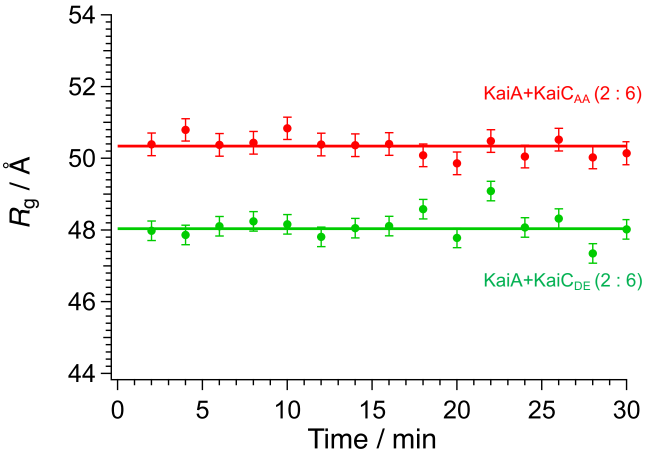


**Supplementary Figure 3. Time dependence of gyration radius.** Red and green circles represent the time dependences of gyration radius ($R_g$) for KaiA+KaiC$_{AA}$ and KaiA+KaiC$_{DE}$ solutions at $x = 2$, respectively. Solid red and green lines mean the fitting curves with linear function for eye guide.

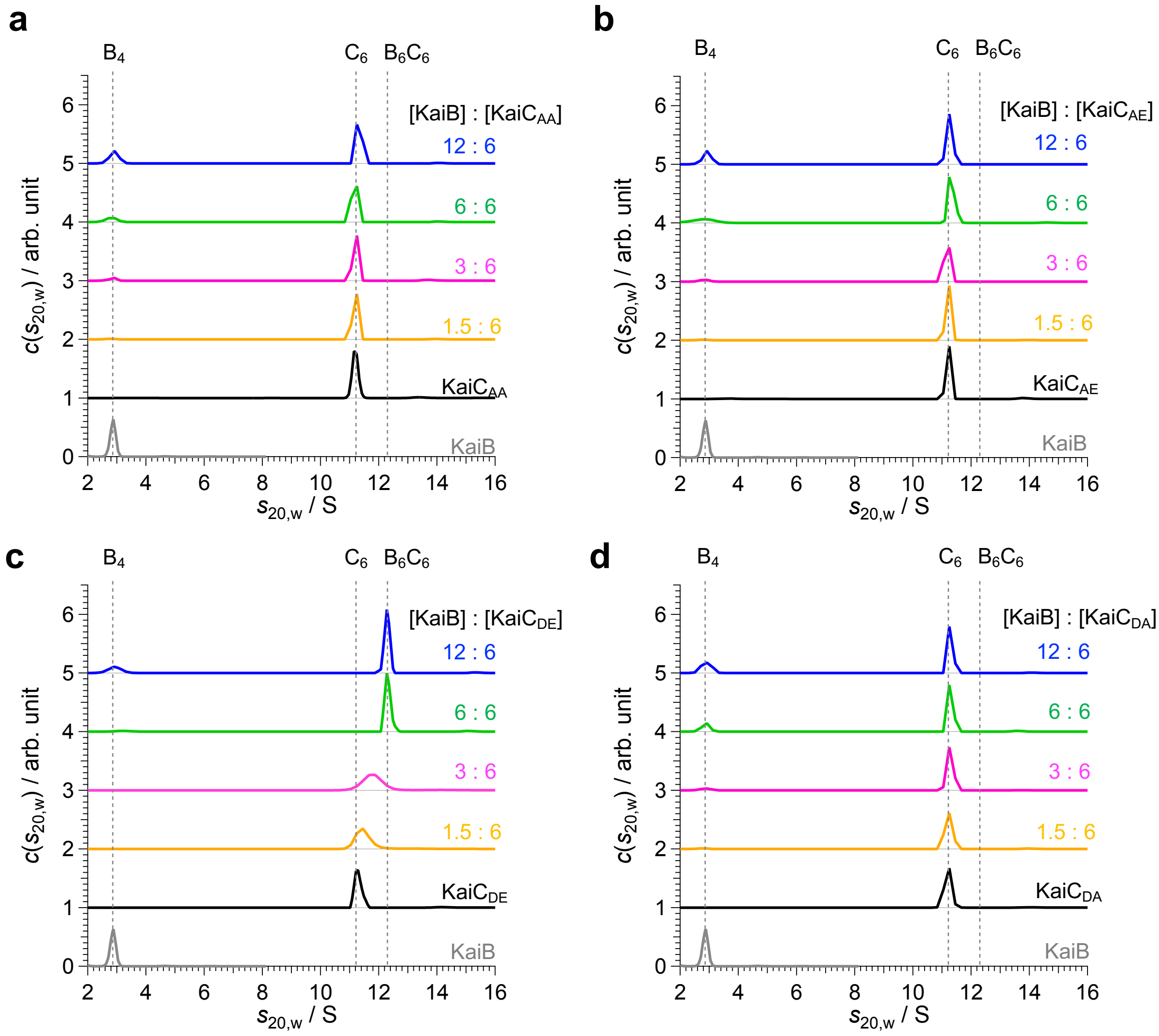


**Supplementary Figure 4. AUC-derived weight concentration distributions from titration of KaiB against KaiC.** Weight-concentration distributions $c(s_{20,w})$ obtained by AUC for KaiB, KaiC, and mix solutions of KaiB+KaiC with various molar mixing ratios. Concentration of KaiC was fixed at 1.0 mg/mL. Panels **a**, **b**, **c**, and **d** show the results for KaiC$_{AA}$, KaiC$_{AE}$, KaiC$_{DE}$, and KaiC$_{DE}$, respectively.

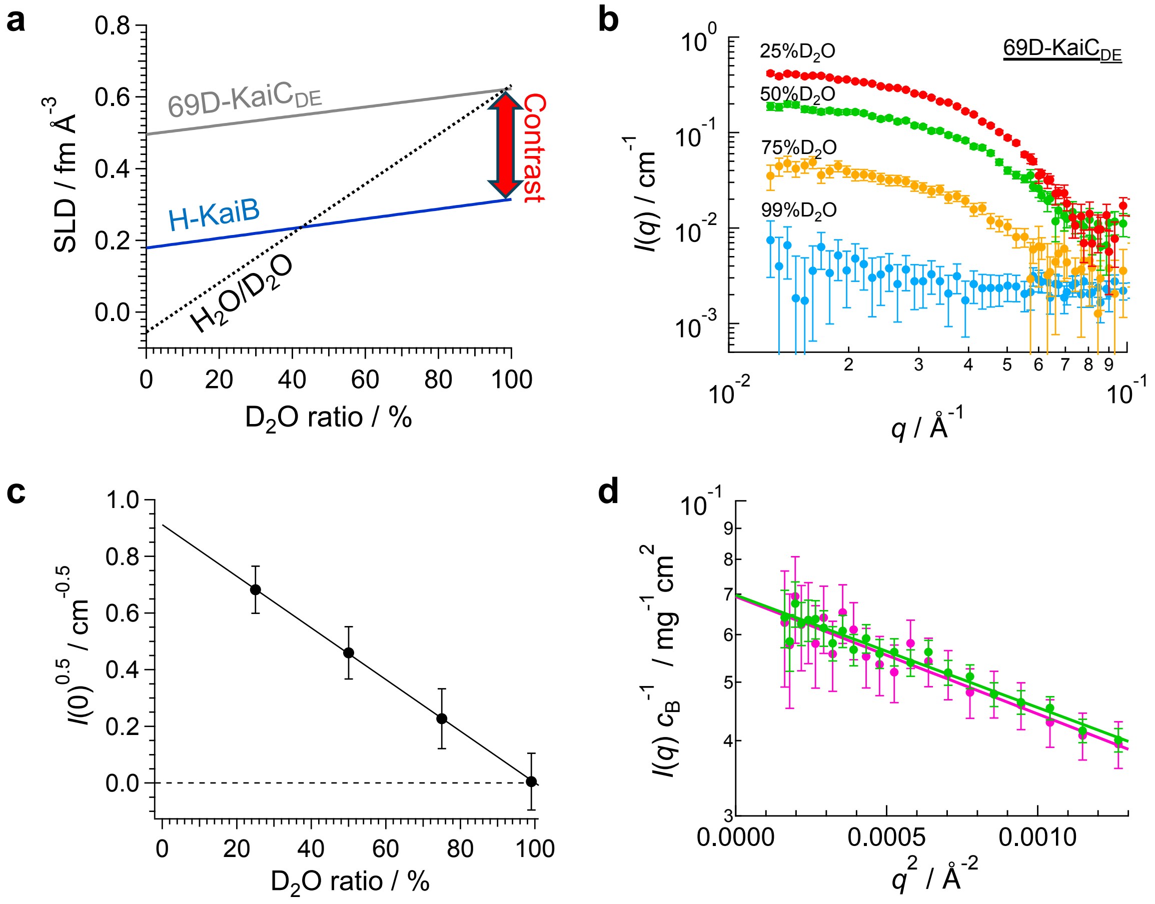


**Supplementary Figure 5. Inversed contrast matching (iCM)-SANS for BC complex. a.** Neutron scattering density map of hydrogenated (H)-KaiB (solid blue line), 69% deuterated (69D)-KaiC$_{DE}$ (solid gray line), and $H_2O/D_2O$ solvent (broken black line) as a function of $D_2O$ ratio. **b.** Scattering intensities $I(q)$ of 69D-KaiC$_{DE}$ in 25%$D_2O$ (red), 50%$D_2O$ (green), 75%$D_2O$ (yellow), and 99%$D_2O$ (cyan). **c.** Square root of forward scattering intensity $I(0)$ , which is proportional to scattering contrast, of 69D-KaiC$_{DE}$ depending on $D_2O$ ratio. From **b** and **c**, invisibility of 69D-KaiC in 99%D2O was validated. **d.** Guinier plots of iCM-SANS profiles for the mix solutions of H-KaiB and 69D-KaiC$_{DE}$ in 99%$D_2O$ buffer at $y$ = 3 (magenta circles) and $y$ = 6 (green circles) of mixing ratios ($y$ = 6[KaiB]/[KaiC]). Scattering intensities were normalized by the weight concentration of KaiB, $c_B$. Solid lines represent the least square fitting with Guinier formula. Forward scattering intensities and gyration radii were summarized in Supplementary Table 2.

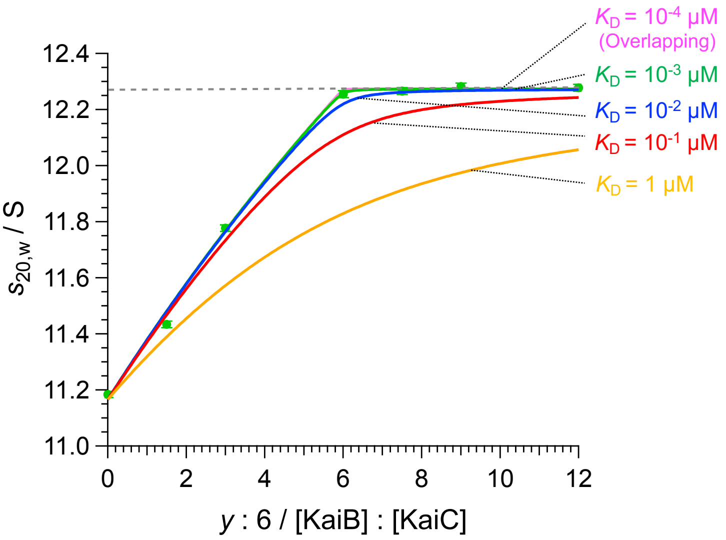


**Supplementary Figure 6. Mixing ratio-dependence of the $s_{20,w}$-value of the peak corresponding to $C_6$/$B_6C_6$ component and calculated $s_{av}$ with various $K_D$.** Green circles represent the experimental results for KaiB+KaiC$_{DE}$ with various mixing ratios of KaiB to KaiC$_{DE}$, [KaiB] : [KaiC$_{DE}$] = $y$ : 6 ($y$ = 0 – 12). Red, yellow, blue, green, and magenta lines are the calculated $s_{av}$ at $K_D$ = 1, $10^{-1}$, $10^{-2}$, $10^{-3}$, and $10^{-4}$ μM, respectively, by the eq. S1 in Supplementary Note 1.

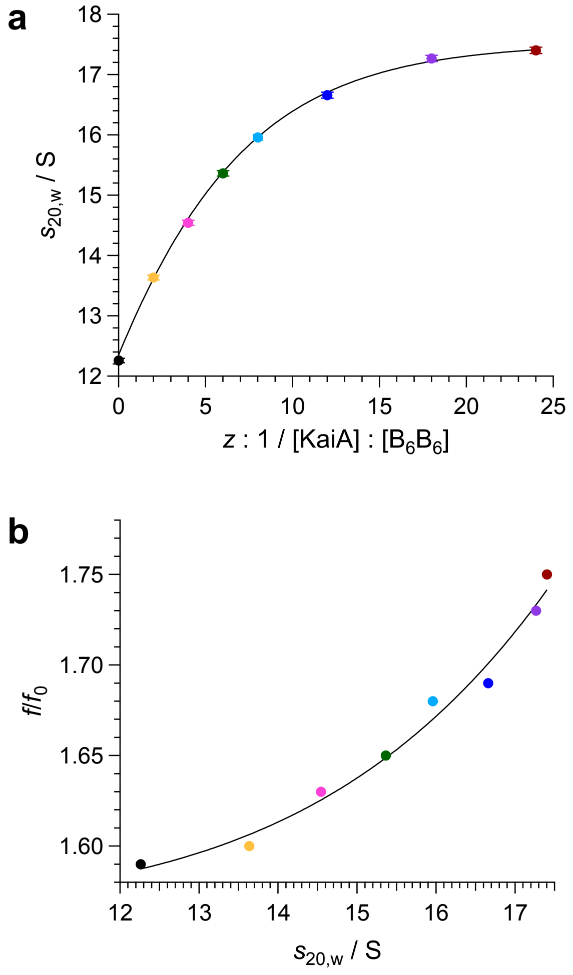


**Supplementary Figure 7. AUC results for ABC complex. a.** Sedimentation coefficient $s_{20,w}$ of ABC complex depending on the mixing ratio $z$ ([KaiA] : [$B_6C_6$] = $z$ : 1). Solid line shows fitting curve with polynomial function for eye-guide. **b.** Relationship between $s_{20,w}$ and frictional ratio $f/f_0$. Solid line shows fitting curve with exponential function for eye-guide. In both panels, the color of circles corresponds to the mixing ratio defined in the Fig.3**a** in the main manuscript: $z$ = 0 (black), 2 (yellow), 4 (magenta), 6 (green), 8 (cyan), 12 (blue), 18 (purple), and 24 (brown).

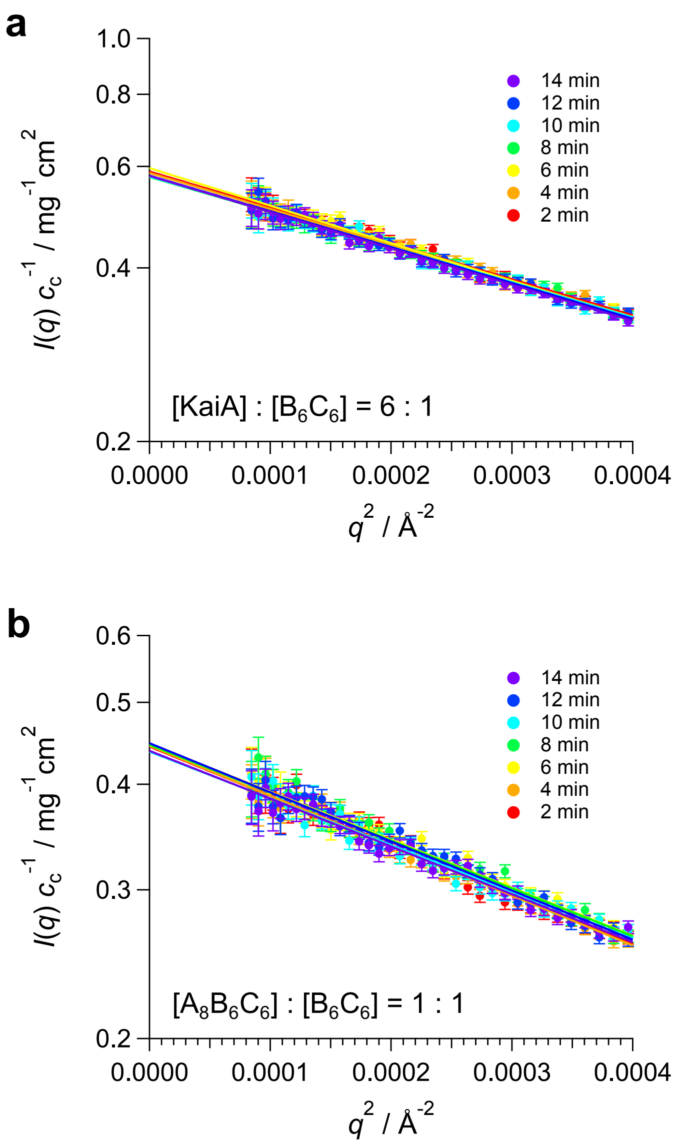


**Supplementary Figure 8. Time resolved SAXS measurements for clock protein complex formation. a.** Guinier plot for time resolved-SAXS profile for the KaiA+$B_6C_6$ solution at the mixing ratio of [KaiA] : [$B_6C_6$] = 6 : 1. **b.** Guinier plot of the time resolved-SAXS profile for the mix solution of $B_6C_6$ and $A_8B_6C_6$. Scattering intensities $I(q)$ are normalized by the weight concentration of KaiC, $c_c$. Solid lines mean the least square fitting with Guinier formula.

**Supplementary Tables:**

**Supplementary Table 1.** Molecular masses and sedimentation coefficients of Kai-protein oligomers and complexes.

| Oligomer /Complex | $B_4$ | $A_2$ | $C_6$ | $A_2C_6$ | $B_6C_6$ | $A_nB_6C_6$ | | | | | |
|---|---|---|---|---|---|---|---|---|---|---|---|
| | | | | | | 2 | 4 | 6 | 8 | 10 | 12 |
| $M_w$ / kDa | 46 | 65 | 348 | 413 | 417 | 481 | 547 | 612 | 678 | 743 | 808 |
| $s_{20,w}$ / S | 2.93 | 3.56 | 11.2 | 12.3 | 12.3 | 13.3 | 14.4 | 15.4 | 16.2 | 16.8 | 17.4 |

**Supplementary Table 2.** KaiC phosphorylation state-dependence of the dissociation constants under the association-dissociation equilibrium of $A_2+C_6 \rightleftarrows A_2C_6$.

| | AA | AE | DE | DA |
|---|---|---|---|---|
| $K_D$ / μM | 0.8 ± 0.1 | 1.3 ± 0.2 | 10.8 ± 0.9 | 8.6 ± 0.8 |

**Supplementary Table 3.** Concentration normalized forward scattering intensities $I(0)c^{-1}$ and gyration radii $R_g$ of KaiA ($A_2$), KaiB ($B_4$), KaiC ($C_6$), and BC complex ($B_6C_6$) obtained by Guinier analysis for SAXS measurements (Supplementary Figure 1).

| | $A_2$ | $B_4$ | $C_6$ | $B_6C_6$ |
|---|---|---|---|---|
| $I(0)c^{-1}$ / mg$^{-1}$ cm$^2$ | 0.0322 ± 0.0001 | 0.0232 ± 0.0002 | 0.176 ± 0.001 | 0.238 ± 0.002* |
| $R_g$ / Å | 29.8 ± 0.2 | 24.7 ± 0.2 | 44.9 ± 0.4 | 48.4 ± 0.4 |

* Normalized by the weight concentration of KaiC ($c_c$).

**Supplementary Table 4.** Forward scattering intensities normalized by weight concentration of KaiB $I(0)c_B^{-1}$ and gyration radii $R_g$ which were obtained by Guinier analysis for iCM-SANS measurements (Supplementary Figure 2**d**). Calculation values were derived from the reference structures of $B_6$ in $B_6C_6$, $B_3$ in $B_3C_6$(i), $B_3$ in $B_3C_6$(ii), and $B_3$ in $B_3C_6$(iii) were based on PDB code 5n8y.

Experimental

| | $y = 3$ | $y = 6$ |
|---|---|---|
| $I(0)c_B^{-1}$ / mg$^{-1}$cm$^2$ | 0.069 ± 0.007 | 0.070 ± 0.003 |
| $R_g$ / Å | 36.6 ± 2.7 | 35.9 ± 1.5 |

Calculation

| | $B_6C_6$ | $B_3C_6$(i) | $B_3C_6$(ii) | $B_3C_6$(iii) | $B_4$ |
|---|---|---|---|---|---|
| $I(0)c_B^{-1}$ / mg$^{-1}$cm$^2$ | 0.072 | 0.036 | 0.036 | 0.036 | 0.047 |
| $R_g$ / Å | 34.2 | 23.6 | 28.1 | 30.1 | 21.3 |
